# Automated segmentation and description of the internal morphology of human permanent teeth by means of micro-CT

**DOI:** 10.1101/2020.10.27.356998

**Authors:** David Haberthür, Ruslan Hlushchuk, Thomas Gerhard Wolf

**Affiliations:** Institute of Anatomy, University of Bern, Switzerland; Department of Restorative, Preventive and Pediatric Dentistry, School of Dental Medicine, University of Bern, Switzerland; Department of Periodontology and Operative Dentistry, University Medical Center of the Johannes-Gutenberg-University Mainz, Mainz, Germany

**Author notes:** These authors contributed equally to this work (Ruslan Hlushchuk, Thomas Gerhard Wolf). Corresponding author; Correspondence preferred via GitHub Issues. Otherwise, send a message to.

**Keywords:** Automated segmentation, Biomedical image analysis, Internal tooth morphology, Micro-CT, Physiological foramen geometry, Root canal configuration

## Abstract

Micro-CT is a powerful tool to analyze and visualize the internal morphology of human permanent teeth. It is increasingly used for investigation of epidemiological questions to provide the dentist with the necessary information required for successful endodontic treatment. The aim of the present paper was to propose an image processing method to automate parts of the work needed to fully describe the internal morphology of human permanent teeth.

One hundred and four human teeth were scanned on a high-resolution micro-CT using an automatic specimen changer. Python code in a Jupyter notebook was used to verify and process the scans, prepare the datasets for description of the internal morphology and to measure the apical region of he tooth.

The presented method offers an easy, non-destructive, rapid and efficient approach to scan, check and preview micro-computer tomographic datasets of a large number of teeth. It is a helpful tool for the detailed description and characterization of the internal morphology of human permanent teeth using automated segmentation by means of micro-CT with full reproducibility and high standardization.

## Introduction

Successful endodontic treatments require a precise knowledge of the external and internal morphology of the teeth [1]. For both surgical and non-surgical interventions it is necessary to know both the complex three-dimensional root canal system with its configurations as well as the details of the apical region of the tooth. This knowledge is necessary to select the correct instruments and materials, thus aiding in important treatment decisions. It also helps to avoid errors that can occur during various steps of clinical endodontic treatment, such as preparation of the access cavity, rinsing, shaping and filling of the root canal system [2]. For example, these errors can include perforations during trepanation or failure in preparing the root canals. A detailed description and understanding of the root canal system is, therefore, essential for the clinical practitioner. At present, there are numerous imaging methods for the morphological description of teeth presented in the literature, including the clearing technique [3], optical microscopy [4], two-dimensional radiography, scanning electron microscopy, or three-dimensional imaging techniques such as cone beam computer tomography and micro-computed tomography (micro-CT) [5].

Micro-CT imaging is a method to non-destructively study objects of interest, namely biomedical samples at high resolution, i.e. in the micrometer range. Micro-CT imaging is well suited for the threedimensional (3D) investigation of teeth since it needs no specialized sample preparation in contrast to what is often needed to image soft tissue samples [6–8]. Combined with software rendering, it is a non-destructive, high-resolution, 3D imaging technique that can precisely depict small morphological structures (< 20 μm) in teeth thus making it superior to other *ex-vivo* methods and therefore, suggested as a gold standard in the field [9–11]. Micro-CT is increasingly used for investigation of epidemiological questions to provide the dentist with necessary information that is a prerequisite for successful endodontic treatment [12–14].

Clinically relevant for the dentist are both the root canal configuration and the physiological procedure. Both parameters are important, as they give information about the expected anatomical conditions and the size of the physiological foramen for clinical purposes. Due to the relatively low cost and batch-scanning capabilities of recent desktop micro-CT systems large cohorts of teeth can be scanned in a relatively short time, generating terabytes of raw data. Such a large amount of data necessitates an efficient, reproducible and automated framework to analyze such large tomographic datasets. Previous works have already analyzed large batches of teeth, but only for a small region of each tooth [15,16] and with a considerable degree of manual input required for tooth segmentation [12,17]. The hereby presented protocol provides an automated segmentation method for *ex vivo* research on extracted human teeth using a four-digit root canal configuration code as well as detection and measurement of the physiological foramen parameters. We achieve this by using free and open-source software [18], considerably increasing the impact and availability of our method for collaborators and other users in the field.

## Materials & Methods

### Tooth selection

A total of 104 extracted human permanent mandibular canines were collected from university medical centers in southwest Germany and Switzerland. All teeth included were extracted for reasons unrelated to the study and are so-called excess material making an institutional review board approval unnecessary for the purpose of this investigation. The teeth were single-rooted and investigated according to their morphologic criteria. Inclusion criteria for teeth selection were complete coronal and root development and the absence of root fracture and resorption, coronal and radicular caries and endodontic treatment. Calculus as well as hard and soft tissue was removed using an ultrasonic scaler. Afterwards, the teeth were placed for one hour in a 3% hydrogen peroxide ultrasonic bath and then stored in 70% ethanol [5,12,15,17,19].

### Micro-CT-based morphological analysis

The 104 samples were imaged on a Bruker SkyScan 1272 high-resolution micro-CT machine (Control software version 1.1.19, Bruker microCT, Kontich, Belgium). To facilitate the scanning of this large batch of samples, we used the automatic sample changer to enable us to scan batches of 16 teeth without any intervention. In addition to the sample changer, the machine is equipped with a Hamamatsu L11871_20 X-ray source and a XIMEA xiRAY16 camera. We used a custom-made sample-holder to scan the teeth on the sample changer. The sample holder was 3D-printed on a Form 2 desktop stereolithography printer (Formlabs, Somerville, Massachusetts, USA) and is freely available online (https://git.io/JJbAZ) as part of a library of sample holders [20]. The X-ray source was set to a tube voltage of 80.0 kV and a tube current of 125.0 μA, the x-ray spectrum was filtered by 1mm of Aluminium prior to incidence onto the sample. For each sample, depending on the sample height, we recorded a set of either 4 or 5 stacked scans overlapping its height. Each stack was recorded with 482 TIFF projections of 1632 × 1092 pixels at every 0.4° over a 180° sample rotation. Every single projection was exposed for 950 ms and 3 projections were averaged to greatly reduce image noise. This resulted in a scan time of approximately 40 minutes per stack and between 2 hours and 40 minutes to 3 hours and 15 minutes per sample. In total, we thus scanned for approximately 13 days. On average, we recorded 7.88 GB of raw data for each tooth, totalling 819 GB for all 104 teeth. The obtained projection images were subsequently reconstructed into a 3D stack of axial PNG images spanning the whole length of each tooth with NRecon (Version 1.7.4.6, Bruker microCT, Kontich Belgium) using a ring artifact correction of 14. The whole process resulted in datasets with an isometric voxel size of 10.0 μm. The teeth were all slightly different in height and on average we had about 2700 reconstructions per teeth and a total of approximately 280000 files for all teeth. The reconstructed PNG slices per tooth are on average 3.13 GB in size, totalling approximately 326 GB for all 104 teeth.

### Image processing

We wrote a Jupyter [21] notebook with Python code which allowed for scans to be checked as soon as they were reconstructed during scanning of the first items in the batch. Re-runs of the notebook added newly scanned and reconstructed teeth to the analysis, facilitating preliminary checks and analysis of already scanned teeth. The notebook used for the analysis presented in this manuscript is freely available online [22]. The important steps of the analysis steps are described in detail below.

### Preparation

In a first step we extracted all necessary parameters from the log file of each scan to store into a *Pandas* [23] dataframe for comparison and verification of all the necessary scan parameters of each scanned tooth with all the others. Afterwards, the preview image of each scan was loaded and an overview image of all the scans was generated (see Figure 1, in which we show a randomly selected subset of the whole tooth cohort).

**Figure 1:**
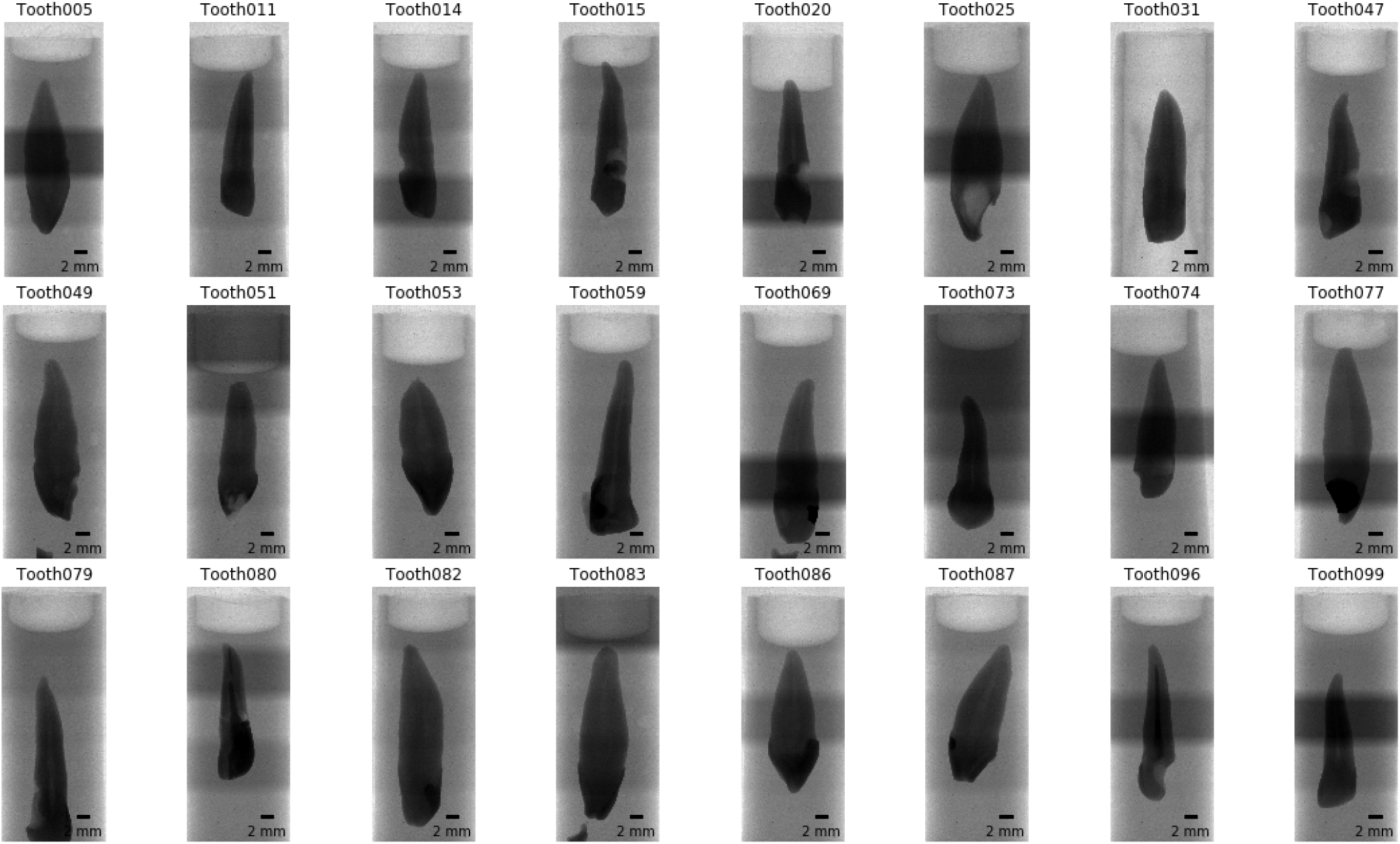
Overview images for a random selection of 24 of the 104 teeth. It is immediately visible that several teeth slipped down in the holder. Since we were particularly interested in the bottom part of the teeth (top in this view) this poses no problem for further analysis. The irregular illumination stems from the rudimentary stitching process of the overview images and is not visible in the reconstructed slices.

We used *Dask* [24] to read the set of axial reconstruction PNG slices for each sample. The *Zarr* library [25] was used to efficiently store a chunked, compressed, two-dimensional array representation of the reconstructions on disk for further analysis, with a total size of 330 GB on disk.

### Dataset cropping

To reduce the size of the data on disk, we cropped the datasets to their minimal amount, i.e. to the smallest cuboid encompassing the full tooth. This was done by segmenting the dataset into tooth and background using a common, fixed gray-value threshold for all datasets. From this refined dataset, we removed small noise by removing small speckles with the remove_small_object function of *scikit-image* [26] and subsequently isolated the biggest object with the find_objects’ function of *SciPy* [27]. The border of this largest object represents the smallest possible region in which the tooth is contained. By removing the empty parts of each three-dimensional dataset containing no information about the teeth, we reduced the size of all datasets nearly three-fold, to approximately 1.11 GB per sample, or a total of 115 GB for all 104 teeth thus facilitating further handling of the data.

### Overview images for visual examination

For quick visual assessment of each of the tooth scans, we extracted overview images for each tooth. After cropping the datasets, we extracted the middle slices and generated the maximum intensity projection (MIP) for each of the anatomical planes [28]. Since the teeth were scanned rotationally invariant, the two anatomical planes along the long axis of the tooth (coronal and sagittal) are not related to the real tooth anatomy and simply correspond to the respective direction in the tomographic dataset.

### Detection of enamel-dentin border

To facilitate the global characterization of the tooth, we extracted slices at four defined locations along the tooth. These slices, located at the EDB, the bottom of the tooth and equidistantly between, were then used to describe the root canal configuration with a 4-digit system and to assess the number of main foramina, both according to a previously proposed method [17]. The location of extracted slices is shown in Figure 2.

**Figure 2:**
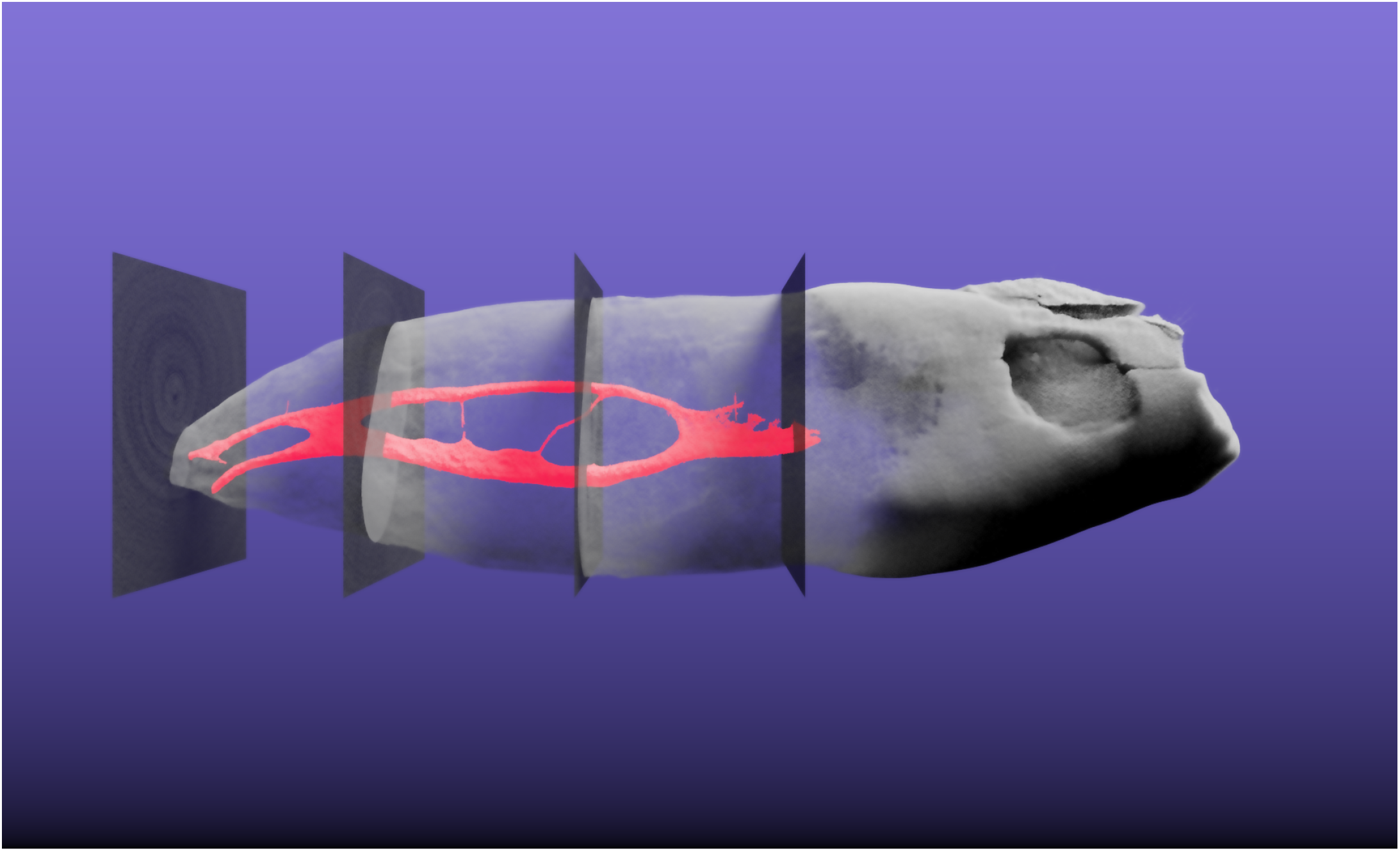
Three-dimensional visualization of tooth sample 045. This tooth is interesting as it features a 1 −2-2/2 root canal configuration as defined by Briseño et al. [17]. The extracted pulpa in shown in red, the tooth itself is shown semitransparently. The four slices which were automatically extracted based on the enamel-dentin border are also visualized semitransparently in their correct 3D position. The whole tooth has a length of 2.39 cm. [A video of the 3D visualization is found in the supplementary materials](https://github.com/habi/zmk-tooth-cohort-method-manuscript/blob/master/content/images/tooth045.mp4?raw=true) of this manuscript.

This was done by calculating the brightness value along the longest axis of the tooth followed by smoothing of the curve using a [locally weighted scatterplot smoothing](https://www.statsmodels.org/devel/generated/statsmodels.nonparametric.smoothers_lowess.lowess.html) implemented in the *statsmodels* library [29]. By finding the maximal derivation of this curve with *NumPy* [30], we were easily able to detect the border between enamel and dentin (EDB). From the part of the tooth below the detected border, we then extracted four equidistant slices. These slices and an overview of the tooth were written to a separate file for each sample, facilitating characterization of each tooth without manually looking for the correct axial reconstruction (see Figure 3).

**Figure 3:**
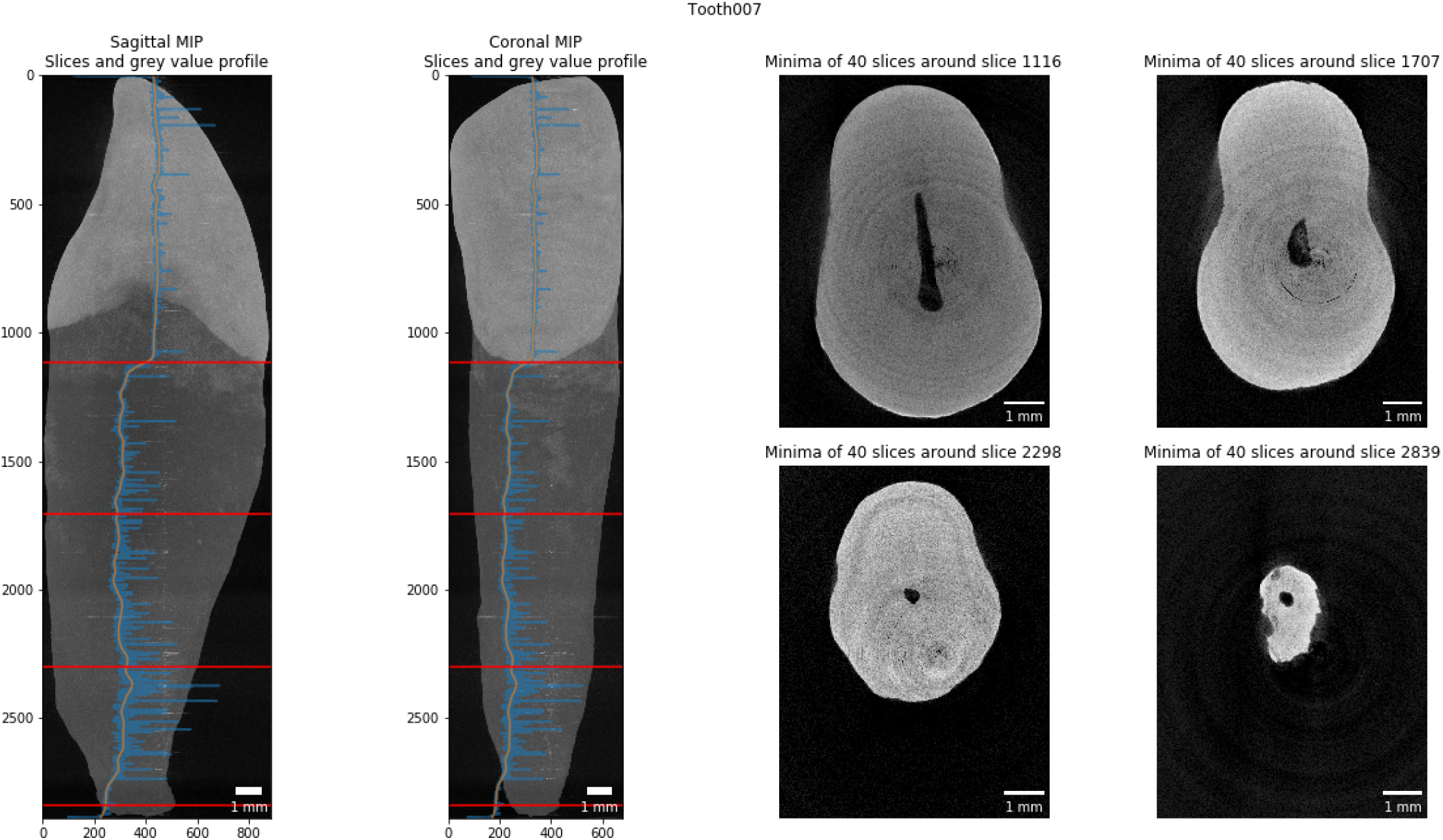
Slice extraction for characterization of a tooth. This tooth features a 1-1-1/1 root canal configuration as defined by Briseño et al. [17]. The blue line in the two leftmost panels shows the gray value plot along the longest tooth axis, the orange line shows the smoothed plot. Based on the largest derivation we detect the enamel-dentin border at slice 1116 of this dataset. Based on the bottom around slice 2839 we extracted the equidistant slices in-between.

### Pulpa extraction

It was of paramount importance for the analysis to visualize the pulpa inside the tooth. We thus wrote a function to extract the pulpa based on its appearance in the axial slices of the datasets. By using an automated Otsu thresholding implemented in *scikit-image* [26] on each slice of the datasets, we separated the tooth from the background. By inversion of the image we select all that is not tooth. From this, we remove all the pixels that touch the image border i.e. the air surrounding the tooth, with the clear_border function. We further removed speckles in the remains and closed small holes in the remaining image data to extract the pulpa from inside the tooth (with the remove_small_objects and remove_small_holes functions, respectively). Datasets of the pulpa have again been written to disk for further analysis and display. Since these are binarized datasets, we were able to store them on disk very efficiently, with the total size of all 104 pulpa datasets being only 309 MB. Display of the datasets for visual assessment was done with *itkwidgets* [31], permitting a basic 3D visualization of each tooth for quality control (an example is shown in Figure 4).

**Figure 4:**
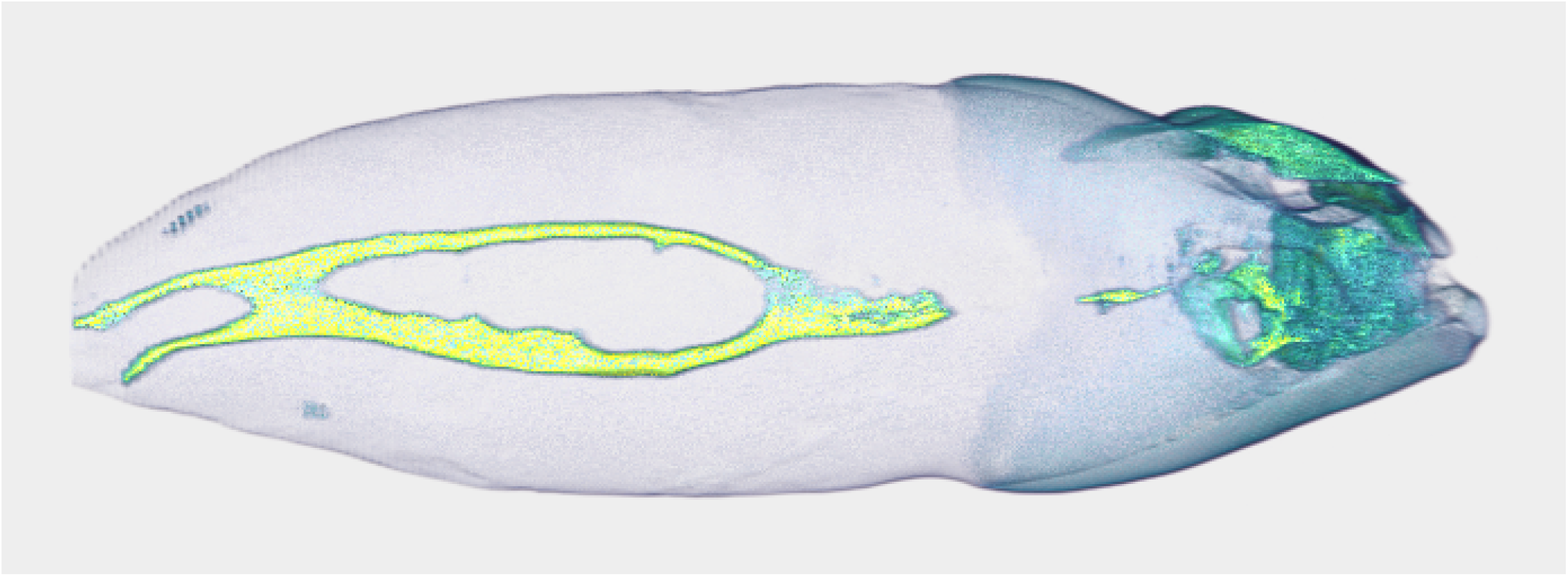
Basic 3D visualization directly from our preparation and analysis pipeline. This tooth is interesting as it features a 1 −2-2/2 root canal configuration as defined by Briseño et al. [17]. The whole tooth has a length of 2.39 cm.

### Analysis of the physiological foramen geometry

The apical foramen of the teeth was evaluated as previously described [19] by assessing the bottom part of each tooth using *Fiji* [32] to scroll through the stack of images and measure the diameter of the physiological and anatomical foramen as well as the distance between the physiological and anatomical foramina. The physiological (main) foramen was defined as one with a diameter of 0.20 mm or more. Foramina with diameters smaller than 0.20 mm were defined as accessory ones [19]. Since we have extracted the pulpa for each tooth, we can easily calculate its diameter at each point, that is, exactly calculate the exact Euclidean distance transform (EDT) where each 3D voxel of the pulpa is labelled with its distance relative to the background (wall of the canal). We used the morphology.distance_transform_edt function of the *SciPy*[27] library for this. To aid the assessment of the geometry of the physiological foramen, we extracted the bottom 3.5 mm part of each tooth, merged the reconstructed slices of the data with the calculated EDT from the pulpa and wrote this data to disk reformatted into sagittal slices. In such a way, the radius of the largest sphere fitted into the pulpa at each point can easily be read off the image upon visual examination. The use of the *Dask* library facilitated efficient reformatting of the datasets and writing them to disk.

## Results and Discussion

High resolution datasets of large batches of teeth were acquired in an efficient manner with minimized operator effort due to the batch-scanning abilities of the desktop micro-CT scanner. The acquired datasets were imaged at a voxel size (10 μm) permitting the analysis of the finest features of interest in the teeth.

The batch-characteristics of the proposed dataset preparation and analysis method makes it easy and efficient to begin processing tooth datasets as scanning of a large batch of teeth is already underway. Our script facilitates short turnaround time for feedback on single scans in the batch, since samples can be processed by the script as soon as they are reconstructed and while other teeth are still being scanned or waiting to be scanned. Cropping the datasets with a simple algorithm - as described in the subsection ‘Dataset cropping’ above - greatly reduces the size of the datasets on disk.

The proposed method is completely devoid of any manual input, all the datasets present on disk are prepared and analyzed automatically. This allows for a highly reproducible and completely unbiased analysis. Previous studies [13,33] have analyzed teeth with a precisely defined manual protocol which necessitated several, accurately performed manual steps, increasing the likelihood of operator error being introduced. This is avoided in the method presented here.

Several teeth contained metal fillings (amalgam) in the crown area, which are difficult to penetrate with the X-ray source available to us. A simple thresholding lead to artefacts that extended to the border of the original dataset, thus for these datasets, there were no gains in disk space. Since the cropping part only influences the Anal size of the dataset on disk and not the extraction of the pulpa from the tooth, it is only of minor concern. Additionally, the implants or metal Allings in several teeth made it impossible to automatically detect the enamel-dentin border. If an implant or Alling is present in the tooth, the largest derivation in the gray value profile along the tooth is situated at the bottom end of the implant. The function to extract the EDB was implemented in a way that a manual extraction of the border and the corresponding slices along the tooth axis was possible.

The reformatting of only the bottom part of the tooth greatly facilitated the analysis of the geometry of the physical foramen of each tooth. While all reconstructions of a single tooth are more than 1 GB in size, these partial datasets are only around 22 MB per tooth. In total, the bottom 3.5 mm of all 104 teeth occupied only 2.3 GB of disk space, making further assessment of the foramen easily and efficiently possible on a standard office laptop.

The non-destructively acquired three-dimensional datasets of the tooth can also be used for additional analysis of tooth morphology, akin to the process outlined by Di Angelo et al. [34], Peters et al. [35] or Paqué et al. [16]. Further work will focus on automatically extracting a description of the physiological foramen which will allow for dentists to gain important information required for a successful root canal treatment.

The hereby presented workflow is based completely on free and open-source software and - can therefore be verified independently by any interested reader. The Jupyter notebook described here is also freely available online [22]. A copy with two samples from the cohort can be run in your browser without installing any software via *Binder*[36] by [clicking a single](https://mybinder.org/v2/gh/habi/zmk-tooth-cohort/master?filepath=ToothAnalysis.ipynb) button in the [README file](https://github.com/habi/zmk-tooth-cohort/blob/master/README.md) of the project repository.

## Conclusions

The presented method offers an efficient approach to scan, check and preview micro-computer tomographic datasets of many teeth. We describe a helpful, free and open-source software tool to prepare datasets for precise description and characterization of the internal morphology of human permanent teeth using automated segmentation of features of interest. Due to the high reproducibility and standardization of the presented method, datasets of large cohorts and populations can be investigated easily and rapidly.

## Acknowledgements

We thank Oleksiy-Zakhar Khoma (Institute of Anatomy, University of Bern, Switzerland) for designing the sample holder that we used for scanning the teeth. We thank Jennifer Fazzari (Institute of Anatomy, University of Bern, Switzerland) for proof-reading the manuscript. We thank Andrea Anderegg and Michael Stiebritz (both Department of Restorative, Preventive and Pediatric Dentistry, School of Dental Medicine, University of Bern, Switzerland) and Valentin Djonov (Institute of Anatomy, University of Bern, Switzerland) for their kind support. This research did not receive any specific grant from funding agencies in the public, commercial, or not-for-profit sectors.

## Contributions

DH scanned the teeth, wrote the analysis notebook and the original draft of the manuscript. RH contributed with discussion during the development of the method and reviewed the manuscript. TGW contributed with ideas and discussion for developing the method, discussed the results, edited and reviewed the manuscript. All authors read and approved the final manuscript.

